## Supplementary Information for "No model to rule them all: a systematic comparison of 83 thermal performance curve models across traits and taxonomic groups"

### Contents

**S1 Models and thermal performance datasets examined in previous studies 2**

**S2 Model fits included in the present study 4**

**S3 Comparison of the ten best-performing models across traits 6**

**S4 Comparison of model performance across datasets of individual enzymes 7**

**S5 TPC descriptor variables 8**

**S6 The topology of the final conditional inference tree 11**

**S7 Thermal performance curve models included in this study 12**

#### S1 Models and thermal performance datasets examined in previous studies

**Supplementary Table 1:** Description of previous studies that compared the performance of alternative TPC models across one or more thermal performance datasets.

| Study | Taxonomic breadth | Trait(s) | Model count | Dataset count |
|---|---|---|---|---|
| Angilletta Jr, M. J. Estimating and comparing thermal performance curves. *J. Therm. Biol.* **31**, 541–545 (2006). | *Sceloporus undulatus* (eastern fence lizard) | maximal sprint speed | 5 | 1 |
| Shi, P. & Ge, F. A comparison of different thermal performance functions describing temperature-dependent development rates. *J. Therm. Biol.* **35**, 225–231 (2010). | *Plutella xylostella* (diamondback moth) and *Bemisia tabaci* (silverleaf whitefly) | development rate | 12 | 2 |
| Krenek, S., Berendonk, T. U. & Petzoldt, T. Thermal performance curves of Paramecium caudatum: a model selection approach. *Eur. J. Protistol.* **47**, 124–137 (2011). | *Paramecium caudatum* (a unicellular protist) | population growth rate | 12 | 4 |
| Shi, P.-J., Reddy, G. V., Chen, L. & Ge, F. Comparison of thermal performance equations in describing temperature-dependent developmental rates of insects: (I) empirical models. *Ann. Entomol. Soc. Am.* **109**, 211–215 (2016). | 9 insect species and *Kampimodromus aberrans* (a mite) | development rate | 6 | 10 |

Supplementary Table 1 – *Continued from previous page*

| | | | | |
|---|---|---|---|---|
| Shi, P.-J., Reddy, G. V., Chen, L. & Ge, F. Comparison of thermal performance equations in describing temperature-dependent developmental rates of insects: (II) two thermodynamic models. *Ann. Entomol. Soc. Am.* **110**, 113–120 (2017). | 9 insect species and *Kampimodromus aberrans* (a mite) | development rate | 2 | 10 |
| Low-Décarie, E. et al. Predictions of response to temperature are contingent on model choice and data quality. *Ecol. Evol.* **7**, 10467–10481 (2017). | mainly phytoplankton and fungi, but also bacteria, ants, and others | mainly population growth rate, but also photosynthesis, filtration rate, and others | 12 | 381 |
| Quinn, B. K. Performance of the SSI development function compared with 33 other functions applied to 79 arthropod species' datasets *J. Therm. Biol.* **102**, 103112 (2021). | arthropods | development rate | 34 | 79 |

#### S2 Model fits included in the present study

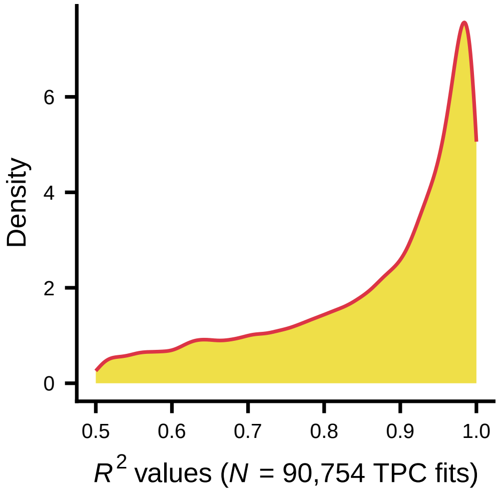


**Supplementary Fig. 1:** $R^2$ values for all TPC model fits that passed our filtering criteria (see Methods in the main text). The distribution is heavily left-skewed, indicating that most model fits that were included in this study were able to accurately represent the underlying measurements of thermal performance.

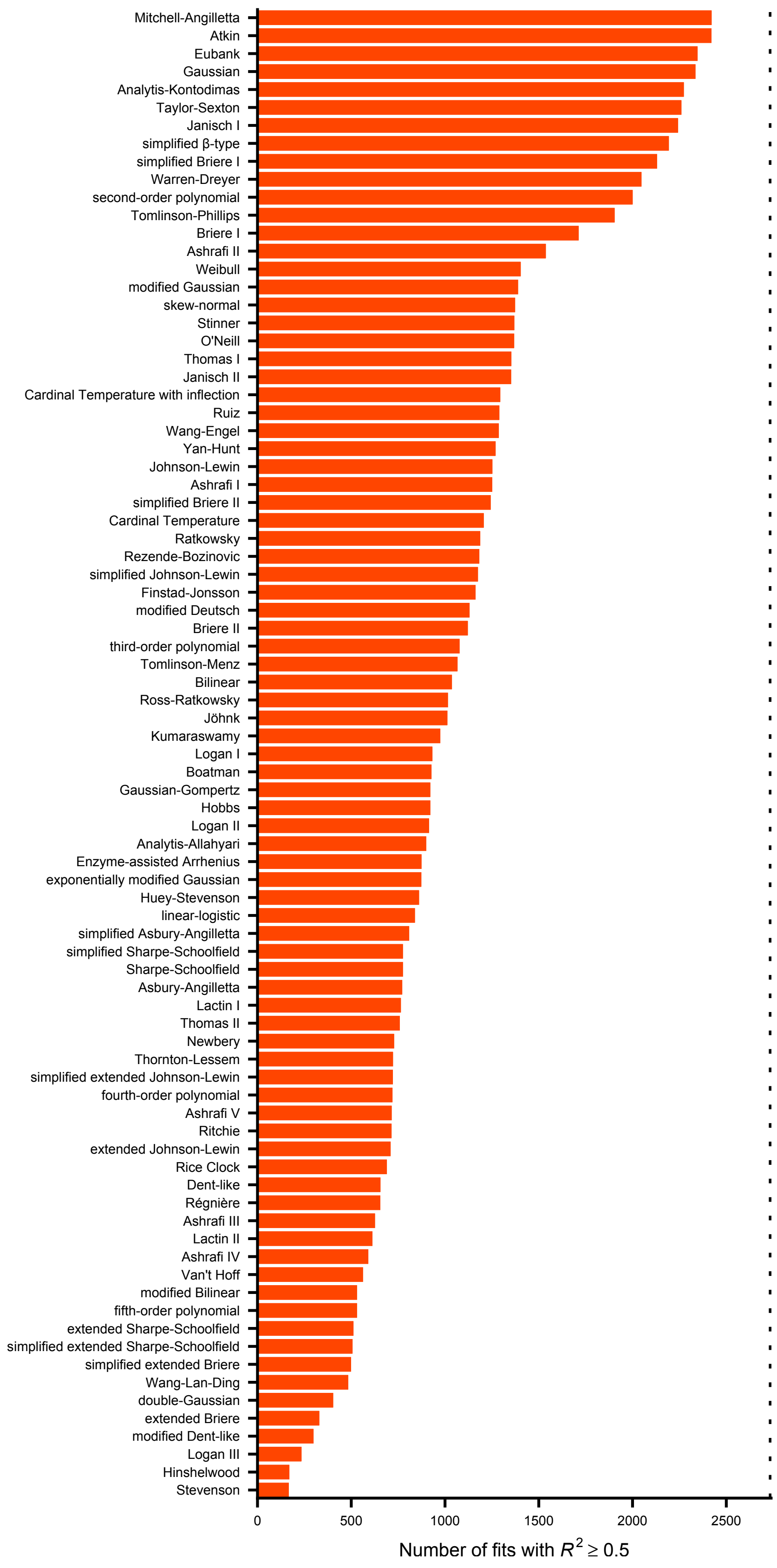


**Supplementary Fig. 2:** The number of fits per model in our study. The dotted line stands for the total number of thermal performance datasets that could be fitted well by at least one model (see Methods in the main text).

#### S3 Comparison of the ten best-performing models across traits

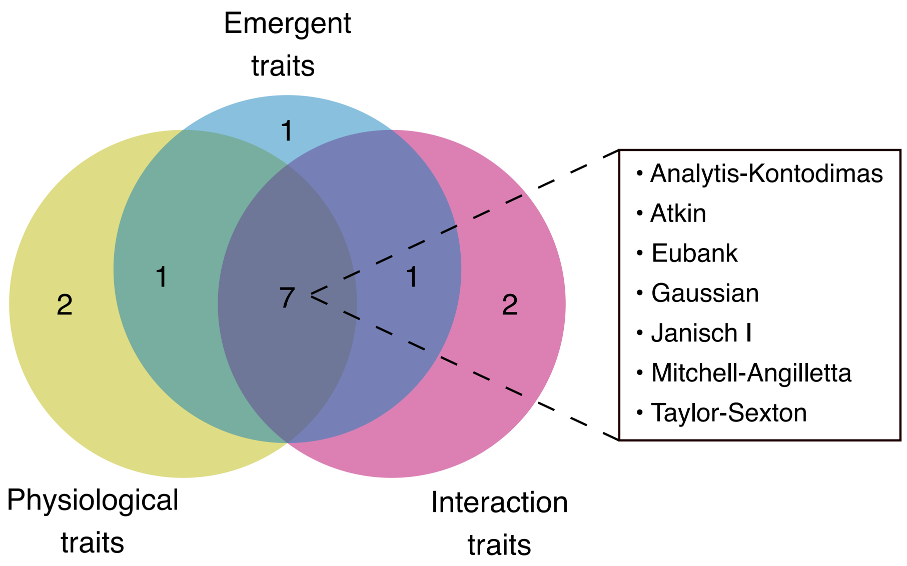


**Supplementary Fig. 3:** Venn diagram of the ten best-performing models (based on their median AICc weights) across thermal performance datasets of physiological, emergent, and interaction traits. The box shows the models that are among the ten best-performing in all three groups.

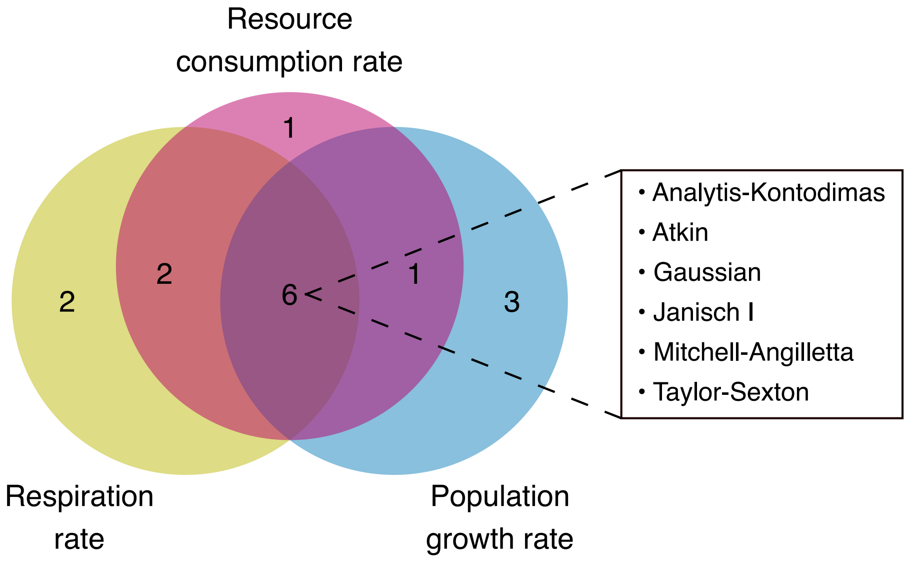


**Supplementary Fig. 4:** Venn diagram of the ten best-performing models (based on their median AICc weights) across thermal performance datasets of respiration rate, population growth rate, and resource consumption rate. The box shows the models that are among the ten best-performing in all three groups.

### S4 Comparison of model performance across datasets of individual enzymes

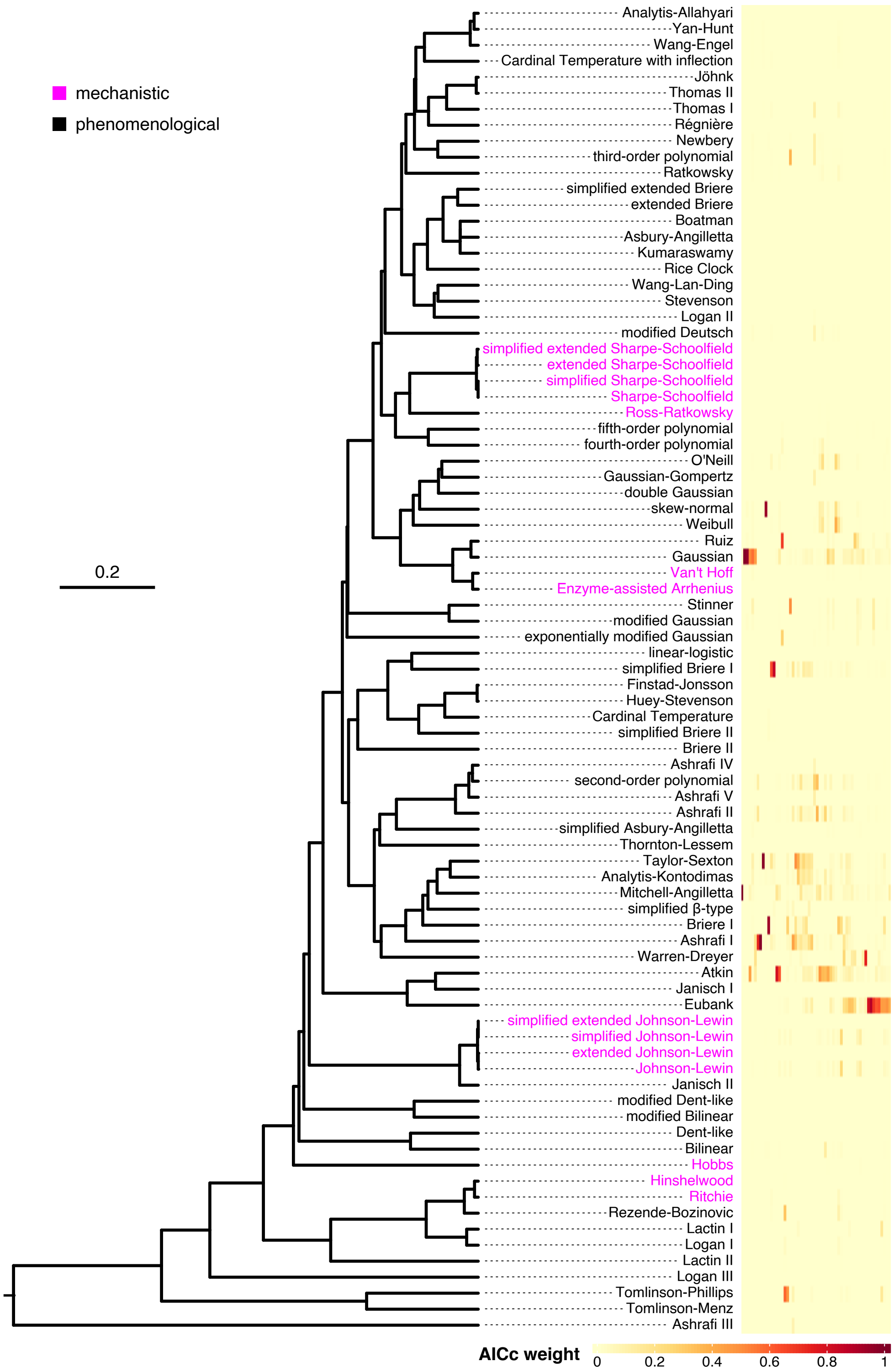


**Supplementary Fig. 5:** Comparison of the performance of our 83 TPC models across 56 thermal performance datasets of individual enzymes. The dendrogram of models was copied from Fig. 3 in the main text.

### S5 TPC descriptor variables

**Supplementary Table 2:** Names and descriptions of the 29 variables used as possible predictors in the multi-output conditional inference regression trees fitted in the present study. The colour of each row stands for variable type. Variables related to sampling resolution are shown in blue, those describing the shape of the TPC in orange, taxonomic variables in grey-green, and trait information variables in pink.

| Variable name | Variable description |
|---|---|
| `n_data_points` | The number of data points across the entire thermal performance dataset. |
| `n_data_points_rise` | As `n_data_points`, but below the thermal optimum. The latter was estimated through a regression of trait measurements against the second order polynomial of temperature.<br><br>We considered a thermal performance dataset to have an estimable thermal optimum if the leading coefficient of the polynomial was negative and the $p$-value of the regression was $< 0.05$.<br><br>If either of these conditions was not met, we estimated the correlation between trait measurements and temperatures. A statistically significant positive/negative correlation would indicate that trait measurements are rising/falling throughout the sampled temperature range.<br><br>If the correlation was not statistically significant, we could not confidently classify the trait measurements as rising, falling, or unimodal and, thus, we assigned NA to this variable. |
| `n_data_points_fall` | As `n_data_points`, but above the thermal optimum. |
| `n_distinct_temperatures` | The number of distinct temperatures across the entire thermal performance dataset. |
| `n_distinct_temperatures_rise` | As `n_distinct_temperatures`, but below the thermal optimum. |

| | |
|---|---|
| n_distinct_temperatures_fall | As n_distinct_temperatures, but above the thermal optimum. |
| correlation_rise | The correlation between trait measurements and temperatures below the thermal optimum. It is a metric of measurement noise. |
| correlation_fall | As correlation_rise, but above the thermal optimum. |
| temperature_range_width | The difference between the maximum and minimum experimental temperatures across the entire thermal performance dataset. |
| temperature_range_width_rise | As temperature_range_width, but below the thermal optimum. |
| temperature_range_width_fall | As temperature_range_width, but above the thermal optimum. |
| median_distance_between_temperatures | The median distance between consecutive temperatures across the entire thermal performance dataset. |
| median_distance_between_temperatures_rise | As median_distance_between_temperatures, but below the thermal optimum. |
| median_distance_between_temperatures_fall | As median_distance_between_temperatures, but above the thermal optimum. |
| median_distance_between_measurements | The median distance between trait measurements at consecutive temperatures across the entire thermal performance dataset, divided by the range of trait measurements. |
| median_distance_between_measurements_rise | As median_distance_between_measurements, but below the thermal optimum. |
| median_distance_between_measurements_fall | As median_distance_between_measurements, but above the thermal optimum. |
| skew_scalar | The skew scalar ($\lambda$) parameter estimate of the skew-normal model (see Supplementary Section S4). Negative values of $\lambda$ correspond to negatively-skewed TPCs, and vice versa. |
| minimum_temperature | The minimum experimental temperature. |

Supplementary Table 2 – *Continued from previous page*

| | |
|---|---|
| maximum_temperature | The maximum experimental temperature. |
| thermal_optimum | See n_data_points_rise. |
| temperature_exponent_rise | Assuming that trait measurements below the thermal optimum increase as a power law of temperature, this variable stands for the exponent of temperature. It is a metric of how steeply the TPC rises up to its peak. |
| temperature_exponent_fall | As temperature_exponent_rise, but above the thermal optimum. |
| minimum_to_maximum_measurement_rise | The ratio of the minimum trait measurement below the thermal optimum over the maximum trait measurement. |
| minimum_to_maximum_measurement_fall | As minimum_to_maximum_measurement_rise, but above the thermal optimum. |
| kingdom | The kingdom to which the organism belongs. |
| phylum | The phylum to which the organism belongs. |
| trait_name | The name of the trait that was measured. |
| trait_group | The type of the trait that was measured (physiological, emergent, or interaction). |

### S6 The topology of the final conditional inference tree

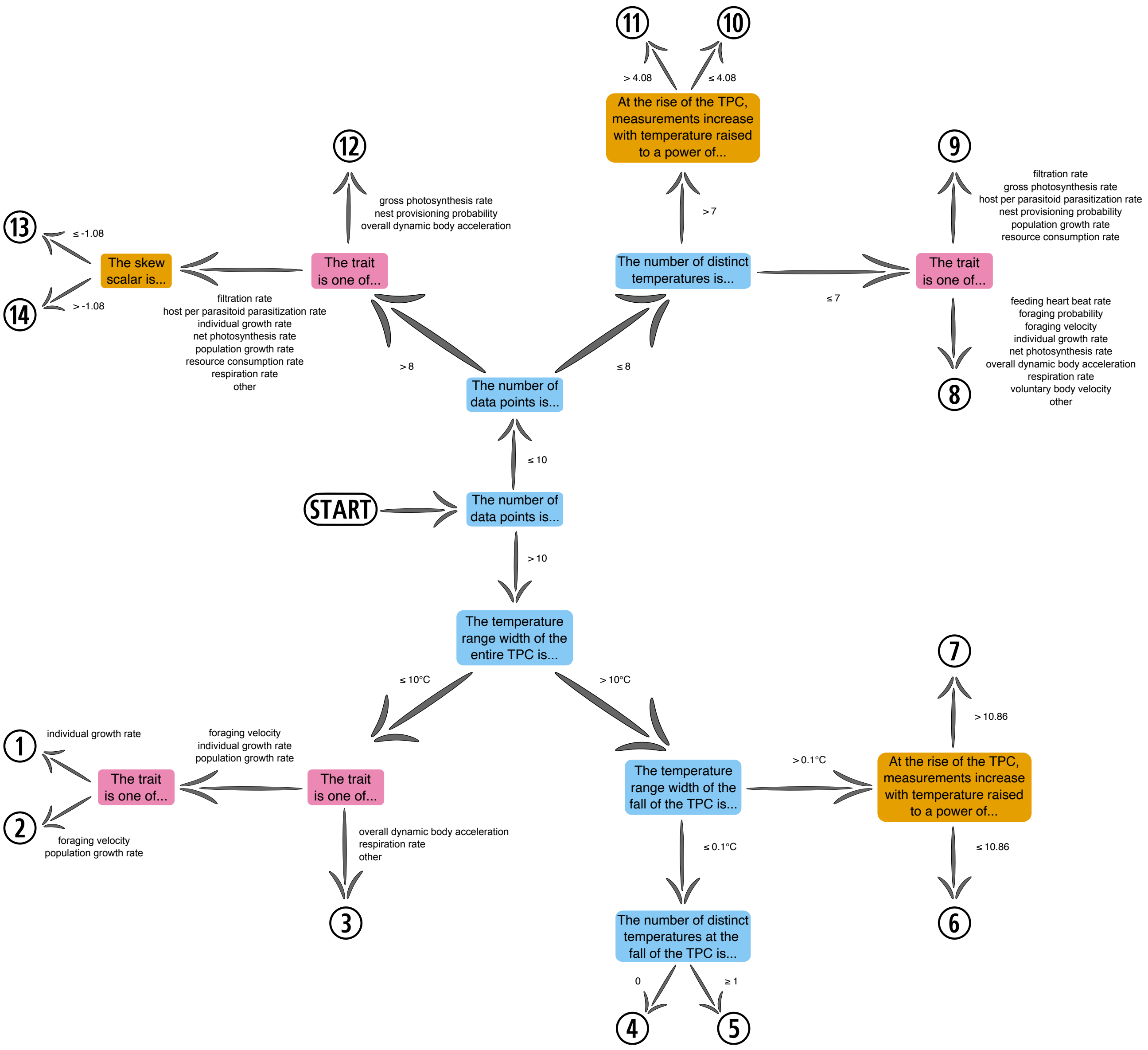


**Supplementary Fig. 6:** The best-fitting conditional inference tree, based on the $R^2$ value obtained across the training data subset. Internal nodes are coloured according to variable type. Blue/orange/pink nodes represent sampling resolution, TPC shape, and trait identity variables, respectively. Circled numbers correspond to the leaf nodes of the tree (i.e., thermal performance datasets with distinct patterns of AICc weights).

#### S7 Thermal performance curve models included in this study

The varying parameters of each model are shown in orange, whereas temperature ($T$), functions, and constants are shown in black.

**1. Analytis-Allahyari (5 parameters)[1]**

$$B(T) = a \cdot \left(\frac{T - T_{\mathrm{min}}}{T_{\mathrm{max}} - T_{\mathrm{min}}}\right)^{b} \cdot \left[1 - \left(\frac{T - T_{\mathrm{min}}}{T_{\mathrm{max}} - T_{\mathrm{min}}}\right)^{d}\right]$$

**2. Analytis-Kontodimas (3 parameters)[2]**

$$B(T) = a \cdot (T - T_{\mathrm{min}})^2 \cdot (T_{\mathrm{max}} - T)$$

**3. Asbury-Angilletta (6 parameters)[3]**

$$B(T) = \psi \cdot \frac{\left(\frac{T-a}{d}\right)^{(c/b)-1} \cdot \left(1 - \frac{T-a}{d}\right)^{[(1-c)/b]-1} \cdot \Gamma\left(\frac{1}{b}\right)}{\Gamma\left(\frac{c}{b}\right) \cdot \Gamma\left(\frac{1-c}{b}\right)} \cdot \exp\left(\frac{-E}{k_{\mathrm{B}} \cdot T}\right)$$

**4. Simplified Asbury-Angilletta (4 parameters)[3]**

$$B(T) = \frac{\left(\frac{T-a}{d}\right)^{(c/b)-1} \cdot \left(1 - \frac{T-a}{d}\right)^{[(1-c)/b]-1} \cdot \Gamma\left(\frac{1}{b}\right)}{\Gamma\left(\frac{c}{b}\right) \cdot \Gamma\left(\frac{1-c}{b}\right)}$$

**5. Ashrafi I (3 parameters)[4]**

$$B(T) = a + b \cdot T^2 \cdot \ln T + c \cdot T^3$$

**6. Ashrafi II (3 parameters)[4]**

$$B(T) = a + b \cdot T^{3/2} + c \cdot T^2$$

**7. Ashrafi III (3 parameters)[4]**

$$B(T) = \frac{1}{a + b \cdot \exp(T) + c \cdot \exp(-T)}$$

**8. Ashrafi IV (4 parameters)[4]**

$$B(T) = a + b \cdot T + c \cdot (\ln T)^2 + d \cdot \sqrt{T}$$

**9. Ashrafi V (4 parameters)[4]**

$$B(T) = a + b \cdot (\ln T)^2 + c \cdot \ln T + \frac{d \cdot \ln T}{T}$$

**10. Atkin (3 parameters)[5]**

$$B(T) = B_0 \cdot (a - b \cdot T)^{T/10}$$

**11. Bilinear (4 parameters)[6]**

$$B(T) = \begin{cases} B_{\text{pk}} \cdot \dfrac{T - T_{\text{min}}}{T_{\text{pk}} - T_{\text{min}}} & \text{for } T_{\text{min}} < T \leq T_{\text{pk}} \\[2ex] B_{\text{pk}} \cdot \dfrac{T_{\text{max}} - T}{T_{\text{max}} - T_{\text{pk}}} & \text{for } T_{\text{pk}} < T < T_{\text{max}} \end{cases}$$

##### 12. Modified bilinear (6 parameters)[7]

$$B(T) = \begin{cases} B_{pk} \cdot \left( \dfrac{T - T_{min}}{T_{pk} - T_{min}} \right)^{a} & \text{for } T_{min} < T \leq T_{pk} \\ \\ B_{pk} \cdot \left( \dfrac{T_{max} - T}{T_{max} - T_{pk}} \right)^{b} & \text{for } T_{pk} < T < T_{max} \end{cases}$$

##### 13. Boatman (5 parameters)[8]

$$B(T) = B_{pk} \cdot \left[ \sin \left( \pi \cdot \left( \frac{T - T_{min}}{T_{max} - T_{min}} \right)^{\theta} \right) \right]^{\Phi}$$

##### 14. Briere I (3 parameters)[9]

$$B(T) = a \cdot T \cdot (T - T_{min}) \cdot \sqrt{T_{max} - T}$$

##### 15. Simplified Briere I (3 parameters)[9]

$$B(T) = a \cdot (T - T_{min}) \cdot \sqrt{T_{max} - T}$$

##### 16. Briere II (4 parameters)[9]

$$B(T) = a \cdot T \cdot (T - T_{min}) \cdot (T_{max} - T)^{1/b}$$

##### 17. Simplified Briere II (4 parameters)[9]

$$B(T) = a \cdot (T - T_{min}) \cdot (T_{max} - T)^{1/b}$$

#### 18. Extended Briere (5 parameters)[9,10]

$$B(T) = a \cdot T \cdot (T - T_{\mathrm{min}})^{b} \cdot (T_{\mathrm{max}} - T)^{c}$$

#### 19. Simplified extended Briere (5 parameters)[9,10]

$$B(T) = a \cdot (T - T_{\mathrm{min}})^{b} \cdot (T_{\mathrm{max}} - T)^{c}$$

#### 20. Cardinal Temperature (4 parameters)[11]

$$B(T) = B_{\mathrm{pk}} \cdot \left(1 - \frac{(T - T_{\mathrm{pk}})^2}{(T - T_{\mathrm{pk}})^2 + T \cdot (T_{\mathrm{max}} + T_{\mathrm{min}} - T) - T_{\mathrm{max}} \cdot T_{\mathrm{min}}}\right)$$

#### 21. Cardinal Temperature with Inflection (4 parameters)[12]

$$B(T) = B_{\mathrm{pk}} \cdot \frac{(T - T_{\mathrm{max}}) \cdot (T - T_{\mathrm{min}})^2}{(T_{\mathrm{pk}} - T_{\mathrm{min}}) \cdot [(T_{\mathrm{pk}} - T_{\mathrm{min}}) \cdot (T - T_{\mathrm{pk}}) - (T_{\mathrm{pk}} - T_{\mathrm{max}}) \cdot (T_{\mathrm{pk}} + T_{\mathrm{min}} - 2 \cdot T)]}$$

#### 22. Dent-like (5 parameters)[13]

$$B(T) = \begin{cases} B_{\mathrm{pk}} \cdot \dfrac{T - T_{\mathrm{min}}}{T_{\mathrm{pk\ (l)}} - T_{\mathrm{min}}} & \text{for } T_{\mathrm{min}} < T < T_{\mathrm{pk\ (l)}} \\ B_{\mathrm{pk}} & \text{for } T_{\mathrm{pk\ (l)}} \leq T \leq T_{\mathrm{pk\ (u)}} \\ B_{\mathrm{pk}} \cdot \dfrac{T_{\mathrm{max}} - T}{T_{\mathrm{max}} - T_{\mathrm{pk\ (u)}}} & \text{for } T_{\mathrm{pk\ (u)}} < T < T_{\mathrm{max}} \end{cases}$$

#### 23. Modified dent-like (7 parameters)[7]

$$B(T) = \begin{cases} B_{\mathrm{pk}} \cdot \left( \dfrac{T - T_{\mathrm{min}}}{T_{\mathrm{pk\ (l)}} - T_{\mathrm{min}}} \right)^{a} & \text{for } T_{\mathrm{min}} < T < T_{\mathrm{pk\ (l)}} \\ B_{\mathrm{pk}} & \text{for } T_{\mathrm{pk\ (l)}} \leq T \leq T_{\mathrm{pk\ (u)}} \\ B_{\mathrm{pk}} \cdot \left( \dfrac{T_{\mathrm{max}} - T}{T_{\mathrm{max}} - T_{\mathrm{pk\ (u)}}} \right)^{b} & \text{for } T_{\mathrm{pk\ (u)}} < T < T_{\mathrm{max}} \end{cases}$$

#### 24. Modified Deutsch (4 parameters)[14,15]

$$B(T) = \begin{cases} B_{\mathrm{pk}} \cdot \exp\left[ - \left( \dfrac{T - T_{\mathrm{pk}}}{2 \cdot \sigma_{\mathrm{p}}} \right)^{2} \right] & \text{for } T \leq T_{\mathrm{pk}} \\ B_{\mathrm{pk}} - B_{\mathrm{pk}} \cdot \left( \dfrac{T - T_{\mathrm{pk}}}{T_{\mathrm{pk}} - T_{\mathrm{max}}} \right)^{2} & \text{for } T > T_{\mathrm{pk}} \end{cases}$$

#### 25. Enzyme-assisted Arrhenius (5 parameters)[16]

$$B(T) = a \cdot \exp\left[ - \frac{E_{\mathrm{b}} - \left( E_{\Delta\mathrm{H}} \cdot \left( 1 - \dfrac{T}{T_{\mathrm{m}}} \right) + E_{\Delta\mathrm{C_p}} \cdot \left( T - T_{\mathrm{m}} - T \cdot \ln \dfrac{T}{T_{\mathrm{m}}} \right) \right)}{k_{\mathrm{B}} \cdot T} \right]$$

#### 26. Eubank (3 parameters)[17]

$$B(T) = \frac{a}{(T - T_{\mathrm{pk}})^{2} + b}$$

#### 27. Finstad-Jonsson (4 parameters)[18]

$$B(T) = a \cdot (T - T_{\mathrm{min}}) \cdot [1 - \exp(b \cdot (T - T_{\mathrm{max}}))]$$

#### 28. Gaussian (3 parameters)[19]

$$B(T) = B_{\mathrm{pk}} \cdot \exp\left[-0.5 \cdot \left(\frac{|T - T_{\mathrm{pk}}|}{a}\right)^2\right]$$

#### 29. Double Gaussian (4 parameters)[20]

$$B(T) = \begin{cases} B_{\mathrm{pk}} \quad \mathrm{e} - \dfrac{-(T - T_{\mathrm{pk}})^2}{2 \cdot a^2} & \text{for } T < T_{\mathrm{pk}} \\ \\ B_{\mathrm{pk}} \quad \mathrm{e} - \dfrac{-(T - T_{\mathrm{pk}})^2}{2 \cdot (a \cdot b)^2} & \text{for } T \geq T_{\mathrm{pk}} \end{cases}$$

#### 30. Modified Gaussian (4 parameters)[19]

$$B(T) = B_{\mathrm{pk}} \cdot \exp\left[-0.5 \cdot \left(\frac{|T - T_{\mathrm{pk}}|}{a}\right)^b\right]$$

#### 31. Exponentially modified Gaussian (5 parameters)[19,21]

$$B(T) = \frac{a \cdot c \cdot \sqrt{2 \cdot \pi}}{2 \cdot d} \cdot \exp\left(\frac{b - T}{d} + \frac{c^2}{2 \cdot d^2}\right) \cdot \left[\frac{d}{|d|} - \mathrm{erf}\left(\frac{b - T}{\sqrt{2} \cdot c} + \frac{c}{\sqrt{2} \cdot d}\right)\right] + f$$

##### 32. Gaussian-Gompertz (5 parameters)[22]

$$B(T) = a \cdot \exp\left[-\exp\left(b \cdot (T - T_{\text{pk}}) - \theta\right) - c \cdot (T - T_{\text{pk}})^2\right]$$

##### 33. Hinshelwood (4 parameters)[23]

$$B(T) = a \cdot \exp\left(\frac{-E_1}{R \cdot T}\right) - b \cdot \exp\left(\frac{-E_2}{R \cdot T}\right)$$

##### 34. Hobbs (4 parameters)[24]

$$B(T) = a \cdot \frac{k_{\text{B}} \cdot T}{h} \cdot \exp\left[-\frac{\Delta H^{\ddagger}_{T_{\text{ref}}} + \Delta C^{\ddagger}_{\text{p}} \cdot (T - T_{\text{ref}})}{R \cdot T} + \frac{\Delta S^{\ddagger}_{T_{\text{ref}}} + \Delta C^{\ddagger}_{\text{p}} \cdot \ln(T/T_{\text{ref}})}{R}\right]$$



##### 35. Huey-Stevenson (5 parameters)[25]

$$B(T) = a \cdot [1 - \exp(-b \cdot (T - T_{\text{min}}))] \cdot [1 - \exp(c \cdot (T - T_{\text{max}}))]$$

##### 36. Janisch I (3 parameters)[26]

$$B(T) = \frac{1}{\frac{m}{2} \cdot \left[a^{T - T_{\text{pk}}} + a^{-(T - T_{\text{pk}})}\right]}$$

##### 37. Janisch II (4 parameters)[26]

$$B(T) = \frac{1}{\frac{m}{2} \cdot \left[a^{T - T_{\text{pk}}} + b^{-(T - T_{\text{pk}})}\right]}$$

**38. Jöhnk (5 parameters)**[27]

$$B(T) = B_{\text{pk}} \cdot \left[1 + a \cdot \left(\left(b^{T-T_{\text{pk}}} - 1\right) - \frac{\ln b}{\ln c} \cdot \left(c^{T-T_{\text{pk}}} - 1\right)\right)\right]$$

**39. Johnson-Lewin (4 parameters)**[28,29]

$$B(T) = B_0 \cdot T \cdot \frac{\exp\left(\frac{-E}{k_{\text{B}}} \cdot \frac{1}{T}\right)}{1 + \frac{E}{E_{\text{D}} - E} \cdot \exp\left[\frac{E_{\text{D}}}{k_{\text{B}}} \cdot \left(\frac{1}{T_{\text{pk}}} - \frac{1}{T}\right)\right]}$$

**40. Extended Johnson-Lewin (5 parameters)**[28–30]

$$B(T) = B_0 \cdot T \cdot \frac{\exp\left(\frac{-E}{k_{\text{B}}} \cdot \frac{1}{T}\right)}{a + \frac{E}{E_{\text{D}} - E} \cdot \exp\left[\frac{E_{\text{D}}}{k_{\text{B}}} \cdot \left(\frac{1}{T_{\text{pk}}} - \frac{1}{T}\right)\right]}$$

**41. Simplified Johnson-Lewin (4 parameters)**[28,29]

$$B(T) = B_0 \cdot \frac{\exp\left(\frac{-E}{k_{\text{B}}} \cdot \frac{1}{T}\right)}{1 + \frac{E}{E_{\text{D}} - E} \cdot \exp\left[\frac{E_{\text{D}}}{k_{\text{B}}} \cdot \left(\frac{1}{T_{\text{pk}}} - \frac{1}{T}\right)\right]}$$

**42. Simplified extended Johnson-Lewin (5 parameters)**[28–30]

$$B(T) = B_0 \cdot \frac{\exp\left(\frac{-E}{k_{\text{B}}} \cdot \frac{1}{T}\right)}{a + \frac{E}{E_{\text{D}} - E} \cdot \exp\left[\frac{E_{\text{D}}}{k_{\text{B}}} \cdot \left(\frac{1}{T_{\text{pk}}} - \frac{1}{T}\right)\right]}$$

**43. Kumaraswamy (5 parameters)[31]**

$$B(T) = a \cdot b \cdot c \cdot \left(\frac{T - T_{\text{min}}}{T_{\text{max}} - T_{\text{min}}}\right)^{b-1} \cdot \left[1 - \left(\frac{T - T_{\text{min}}}{T_{\text{max}} - T_{\text{min}}}\right)^{b}\right]^{c-1}$$

**44. Lactin I (3 parameters)[32]**

$$B(T) = \exp(\rho \cdot T) - \exp\left(\rho \cdot T_{\text{max}} - \frac{T_{\text{max}} - T}{\Delta T}\right)$$

**45. Lactin II (4 parameters)[32]**

$$B(T) = \exp(\rho \cdot T) - \exp\left(\rho \cdot T_{\text{max}} - \frac{T_{\text{max}} - T}{\Delta T}\right) + \lambda$$

**46. Linear-logistic (5 parameters)[33]**

$$B(T) = a \cdot (T - T_{\text{min}}) \cdot \frac{1 - \exp[-b \cdot (T_{\text{max}} - T)]}{1 + \exp[-b \cdot (c - T)]}$$

**47. Logan I (4 parameters)[34]**

$$B(T) = \psi \cdot \left[\exp(\rho \cdot T) - \exp\left(\rho \cdot T_{\text{max}} - \frac{T_{\text{max}} - T}{\Delta T}\right)\right]$$

**48. Logan II (5 parameters)[34]**

$$B(T) = a \cdot \left[\frac{1}{1 + k \cdot \exp(-\rho \cdot T)} - \exp\left(-\frac{T_{\text{max}} - T}{\Delta T}\right)\right]$$

**49. Logan III (4 parameters)**[35]

$$B(T) = \psi \cdot \left[ \frac{T^2}{T^2 + D^2} - \exp\left( - \frac{T_{\max} - T}{\Delta T} \right) \right]$$

**50. Mitchell-Angilletta (3 parameters)**[36]

$$B(T) = \frac{a}{2 \cdot b} \cdot \left[ 1 + \cos\left( \frac{T - T_{\text{pk}}}{b} \cdot \pi \right) \right]$$

**51. Newbery (4 parameters)**[37]

$$B(T) = a + b \cdot T + c \cdot [1 - \exp(d \cdot T^2)]$$



**52. O'Neill (4 parameters)**[38]

$$B(T) = B_{\text{pk}} \cdot V^X \cdot \exp[X \cdot (1 - V)] \quad \textbf{where} \quad V = \frac{T_{\textbf{max}} - T}{T_{\textbf{max}} - T_{\textbf{pk}}}, \quad W = (Q_{\textbf{10}} - 1) \cdot (T_{\textbf{max}} - T_{\textbf{pk}}), \quad \textbf{and} \ X = \frac{W^2 \cdot \left( 1 + \sqrt{1 + \frac{40}{W}} \right)^2}{400}.$$

**53. Second-order polynomial (3 parameters)**

$$B(T) = a + b \cdot T + c \cdot T^2$$

**54. Third-order polynomial (4 parameters)**

$$B(T) = a + b \cdot T + c \cdot T^2 + d \cdot T^3$$

#### 55. Fourth-order polynomial (5 parameters)

$$B(T) = a + b \cdot T + c \cdot T^2 + d \cdot T^3 + f \cdot T^4$$

#### 56. Fifth-order polynomial (6 parameters)

$$B(T) = a + b \cdot T + c \cdot T^2 + d \cdot T^3 + f \cdot T^4 + g \cdot T^5$$

#### 57. Ratkowsky (4 parameters)[39]

$$B(T) = [a \cdot (T - T_{\text{min}}) \cdot (1 - \exp(b \cdot (T - T_{\text{max}})))]^2$$

#### 58. Régnière (6 parameters)[40]

$$B(T) = \psi \cdot \left[\exp(\rho \cdot (T - T_{\text{min}})) - \frac{T_{\text{max}} - T}{T_{\text{max}} - T_{\text{min}}} \cdot \exp\left(-\rho \cdot \frac{T - T_{\text{min}}}{\Delta T_{\text{low}}}\right) - \frac{T - T_{\text{min}}}{T_{\text{max}} - T_{\text{min}}} \cdot \exp\left(\rho \cdot (T_{\text{max}} - T_{\text{min}}) - \frac{T_{\text{max}} - T}{\Delta T_{\text{high}}}\right)\right]$$

#### 59. Rezende-Bozinovic (4 parameters)[41]

$$B(T) = \begin{cases} B_0 \cdot \exp\left(\dfrac{T \cdot \ln Q_{10}}{10}\right) & \text{for } T \leq T_{\text{th}} \\[2ex] B_0 \cdot \exp\left(\dfrac{T \cdot \ln Q_{10}}{10}\right) \cdot [1 - d \cdot (T - T_{\text{th}})^2] & \text{for } T > T_{\text{th}} \end{cases}$$

**60. Rice Clock (6 parameters)**[42]

$$B(T) = \begin{cases} B_{\mathrm{pk}} \cdot \Phi & \text{for } \Phi \leq 1 \\ \\ B_{\mathrm{pk}} & \text{for } \Phi > 1 \end{cases} \quad \textbf{where} \quad \Phi = \left(\frac{T - T_{\mathbf{min}}}{T_{\mathbf{pk}} - T_{\mathbf{min}}}\right)^{a} \cdot \left(\frac{T_{\mathbf{max}} - T}{T_{\mathbf{max}} - T_{\mathbf{pk}}}\right)^{b}$$

**61. Ritchie (4 parameters)**[43]

$$B(T) = R \cdot d_0 \cdot \exp\left(\frac{-E_{\mathrm{D}}}{R \cdot T}\right) \cdot \left(\frac{\Delta E}{R \cdot T} + a\right)$$

**62. Ross-Ratkowsky (5 parameters)**[44]

$$B(T) = \frac{a \cdot T \cdot \exp\left(\dfrac{-\Delta H_{\mathrm{A}}^{\ddagger}}{R \cdot T}\right)}{1 + \exp\left[-n \cdot \dfrac{\Delta H^{*} - 18.1 \cdot T + \Delta C_{\mathrm{p}} \cdot \left(T - 373.6 - T \cdot \ln \dfrac{T}{385.2}\right)}{R \cdot T}\right]}$$

**63. Ruiz (4 parameters)**[45]

$$B(T) = B_0 + \Delta B_{\mathrm{pk}} \cdot \exp[-a \cdot (T - T_{\mathrm{pk}})^2]$$

**64. Sharpe-Schoolfield (6 parameters)**[46]

$$B(T) = \frac{B_0 \cdot \dfrac{T}{T_{\mathrm{ref}}} \cdot \exp\left[\dfrac{\Delta H_{\mathrm{A}}^{\neq}}{R} \cdot \left(\dfrac{1}{T_{\mathrm{ref}}} - \dfrac{1}{T}\right)\right]}{1 + \exp\left[\dfrac{\Delta H_{\mathrm{L}}}{R} \cdot \left(\dfrac{1}{T_{\mathrm{L50}}} - \dfrac{1}{T}\right)\right] + \exp\left[\dfrac{\Delta H_{\mathrm{H}}}{R} \cdot \left(\dfrac{1}{T_{\mathrm{H50}}} - \dfrac{1}{T}\right)\right]}$$

**65. Extended Sharpe-Schoolfield (7 parameters)**[30,46]

$$B(T) = \frac{B_0 \cdot \frac{T}{T_{\text{ref}}} \cdot \exp\left[\frac{\Delta H_{\text{A}}^{\neq}}{R} \cdot \left(\frac{1}{T_{\text{ref}}} - \frac{1}{T}\right)\right]}{a + \exp\left[\frac{\Delta H_{\text{L}}}{R} \cdot \left(\frac{1}{T_{\text{L50}}} - \frac{1}{T}\right)\right] + \exp\left[\frac{\Delta H_{\text{H}}}{R} \cdot \left(\frac{1}{T_{\text{H50}}} - \frac{1}{T}\right)\right]}$$

**66. Simplified Sharpe-Schoolfield (6 parameters)**[46]

$$B(T) = \frac{B_0 \cdot \exp\left[\frac{\Delta H_{\text{A}}^{\neq}}{R} \cdot \left(\frac{1}{T_{\text{ref}}} - \frac{1}{T}\right)\right]}{1 + \exp\left[\frac{\Delta H_{\text{L}}}{R} \cdot \left(\frac{1}{T_{\text{L50}}} - \frac{1}{T}\right)\right] + \exp\left[\frac{\Delta H_{\text{H}}}{R} \cdot \left(\frac{1}{T_{\text{H50}}} - \frac{1}{T}\right)\right]}$$

**67. Simplified extended Sharpe-Schoolfield (7 parameters)**[30,46]

$$B(T) = \frac{B_0 \cdot \exp\left[\frac{\Delta H_{\text{A}}^{\neq}}{R} \cdot \left(\frac{1}{T_{\text{ref}}} - \frac{1}{T}\right)\right]}{a + \exp\left[\frac{\Delta H_{\text{L}}}{R} \cdot \left(\frac{1}{T_{\text{L50}}} - \frac{1}{T}\right)\right] + \exp\left[\frac{\Delta H_{\text{H}}}{R} \cdot \left(\frac{1}{T_{\text{H50}}} - \frac{1}{T}\right)\right]}$$

**68. Simplified $\beta$ type (3 parameters)**[47]

$$B(T) = \rho \cdot \left(a - \frac{T}{10}\right) \cdot \left(\frac{T}{10}\right)^{b}$$

**69. Skew-normal (4 parameters)**[48]

$$B(T) = a \cdot \exp\left[\frac{-(T-z)^2}{\sigma^2}\right] \cdot \left[1 + \text{erf}\left(\frac{\lambda \cdot (T-z)}{\sigma}\right)\right]$$

#### 70. Stevenson (6 parameters)[49]

$$B(T) = a \cdot \frac{1 - \exp[b \cdot (T - T_{\text{max}})]}{1 + c \cdot \exp[-d \cdot (T - T_{\text{min}})]}$$

#### 71. Stinner (4 parameters)[50]

$$B(T) = \begin{cases} B_{\text{pk}} \cdot \dfrac{1 + \exp(a + b \cdot T_{\text{pk}})}{1 + \exp(a + b \cdot T)} & \text{for } T \leq T_{\text{pk}} \\[2ex] B_{\text{pk}} \cdot \dfrac{1 + \exp(a + b \cdot T_{\text{pk}})}{1 + \exp[a + b \cdot (2 \cdot T_{\text{pk}} - T)]} & \text{for } T > T_{\text{pk}} \end{cases}$$

#### 72. Taylor-Sexton (3 parameters)[51]

$$B(T) = B_{\text{pk}} \cdot \frac{-(T - T_{\text{min}})^4 + 2 \cdot (T - T_{\text{min}})^2 \cdot (T_{\text{pk}} - T_{\text{min}})^2}{(T_{\text{pk}} - T_{\text{min}})^4}$$

#### 73. Thomas I (4 parameters)[52]

$$B(T) = a \cdot \exp(b \cdot T) \cdot \left[1 - \left(\frac{T - z}{\frac{w}{2}}\right)^2\right]$$

This model is mathematically equivalent to that fitted by Norberg, 2004[53].

#### 74. Thomas II (5 parameters)[54]

$$B(T) = a \cdot \exp(b \cdot T) - [c + d \cdot \exp(f \cdot T)]$$

#### 75. Thornton-Lessem (6 parameters)[55]

$$B(T) = a \cdot \frac{K_1 \cdot \exp\left[\frac{T - T_1}{T_2 - T_1} \cdot \ln \frac{0.98 \cdot (1 - K_1)}{K_1 \cdot (1 - 0.98)}\right]}{1 + K_1 \cdot \left[\exp\left(\frac{T - T_1}{T_2 - T_1} \cdot \ln \frac{0.98 \cdot (1 - K_1)}{K_1 \cdot (1 - 0.98)}\right) - 1\right]} \cdot \frac{K_4 \cdot \exp\left[\frac{T_3 - T}{T_3 - T_2} \cdot \ln \frac{0.98 \cdot (1 - K_4)}{K_4 \cdot (1 - 0.98)}\right]}{1 + K_4 \cdot \left[\exp\left(\frac{T_3 - T}{T_3 - T_2} \cdot \ln \frac{0.98 \cdot (1 - K_4)}{K_4 \cdot (1 - 0.98)}\right) - 1\right]}$$

#### 76. Tomlinson-Menz (4 parameters)[56]

$$B(T) = a \cdot [\exp(b \cdot T) - \exp(c - T) - \exp(T - d)]$$

#### 77. Tomlinson-Phillips (3 parameters)[57]

$$B(T) = a \cdot [\exp(b \cdot T) - \exp(T - c)]$$

#### 78. Van't Hoff (4 parameters)[58]

$$B(T) = a \cdot \exp\left(\frac{-b}{T}\right) \cdot T^{c} \cdot \exp(d \cdot T)$$

#### 79. Wang-Engel (4 parameters)[59]

$$B(T) = B_{\text{pk}} \cdot \frac{2 \cdot (T - T_{\min})^{a} \cdot (T_{\text{pk}} - T_{\min})^{a} - (T - T_{\min})^{2 \cdot a}}{(T_{\text{pk}} - T_{\min})^{2 \cdot a}}$$

#### 80. Wang-Lan-Ding (7 parameters)[60]

$$B(T) = k \cdot \frac{[1 - \exp(-a \cdot (T - T_{\min}))] \cdot [1 - \exp(b \cdot (T - T_{\max}))]}{1 + \exp[-d \cdot (T - T_0)]}$$

**81. Warren-Dreyer (3 parameters)[61]**

$$B(T) = B_{\mathrm{pk}} \cdot \exp\left[-0.5 \cdot \left(\frac{\ln\dfrac{T}{T_{\mathrm{pk}}}}{a}\right)^2\right]$$

**82. Weibull (4 parameters)[62]**

$$B(T) = B_{\mathrm{pk}} \cdot \left(\frac{c-1}{c}\right)^{(1-c)/c} \cdot \left[\frac{T - T_{\mathrm{pk}}}{b} + \left(\frac{c-1}{c}\right)^{1/c}\right]^{c-1} \cdot \exp\left[-\left(\frac{T - T_{\mathrm{pk}}}{b} + \left(\frac{c-1}{c}\right)^{1/c}\right)^{c} + \frac{c-1}{c}\right]$$

**83. Yan-Hunt (4 parameters)[63]**

$$B(T) = B_{\mathrm{pk}} \cdot \frac{T_{\mathrm{max}} - T}{T_{\mathrm{max}} - T_{\mathrm{pk}}} \cdot \left(\frac{T - T_{\mathrm{min}}}{T_{\mathrm{pk}} - T_{\mathrm{min}}}\right)^{\dfrac{T_{\mathrm{pk}} - T_{\mathrm{min}}}{T_{\mathrm{max}} - T_{\mathrm{pk}}}}$$
